# Improving fold activation of small transcription activating RNAs (STARs) with rational RNA engineering strategies

**DOI:** 10.1101/022772

**Authors:** Sarai Meyer, James Chappell, Sitara Sankar, Rebecca Chew, Julius B. Lucks

## Abstract

Engineered RNAs have become integral components of the synthetic biology and bioengineering toolbox for controlling gene expression. We recently expanded this toolbox by creating small transcription activating RNAs (STARs) that act by disrupting the formation of a target transcriptional terminator hairpin placed upstream of a gene. While STARs are a promising addition to the repertoire of RNA regulators, much work remains to be done to optimize the fold activation of these systems. Here we apply rational RNA engineering strategies to improve the fold activation of two STAR regulators. We demonstrate that a combination of promoter strength tuning and multiple RNA stabilization strategies can improve fold activation from 5.4-fold to 13.4-fold for a STAR regulator derived from the pbuE riboswitch terminator. We then validate the generality of our approach and show that these same strategies improve fold activation from 2.1-fold to 14.6-fold for an unrelated STAR regulator. We also establish that the optimizations preserve the orthogonality of these STARs between themselves and a set of antisense RNA transcriptional repressors, enabling these optimized STARs to be used in more sophisticated circuits. These optimization strategies open the door for creating a generation of additional STARs to use in a broad array of biotechnologies.

## Introduction

Natural and engineered RNA regulators have become powerful components of our toolbox for precisely regulating gene expression (Chappell et al. 2013). This is in large part due to advances in our understanding of RNA biology that have uncovered a vast range of regulatory functions performed by naturally occurring RNAs (Cech and Steitz, 2014; Chappell et al., 2013). Many of these functions involve the regulation of the fundamental processes of gene expression, including mRNA degradation (Collins et al., 2007; Filipowicz et al., 2008; Storz et al., 2011), translation (Gottesman and Storz, 2011; Nou and Kadner, 2000; Winkler et al., 2002), and transcription elongation (Brantl and Wagner, 2000). Recent work has further revealed how these functions are intimately linked to the structure of the regulatory RNAs and the structural rearrangements they induce in their targets (DebRoy et al., 2014). This in turn has enabled significant advances in design approaches that use computational RNA structure prediction algorithms to design synthetic RNAs that adopt specific conformations to perform their regulatory function (Green et al., 2014; Rodrigo et al., 2012). RNA regulators thus represent a versatile and designable platform for controlling gene expression and have been used in a number of recent applications, including the creation of synthetic RNA regulatory gene expression switches (Ceres et al., 2013a; Lynch et al., 2007; Wachsmuth et al., 2013), RNA–only logic gates (Lucks et al., 2011, Chappell et al. 2015), RNA transcriptional networks (Bhadra and Ellington, 2014; Lucks et al., 2011; Takahashi et al., 2014), and RNA-based diagnostics (Pardee et al., 2014).

For bacterial systems, small RNAs (sRNAs) have proven to be particularly well suited to engineering approaches that optimize and alter their function. sRNAs typically act through Watson-Crick base pairing between a sense target RNA, usually located upstream of the gene to be controlled, and a *trans*-acting antisense RNA. By itself, the sense target can fold into structures that block or allow specific aspects of gene expression–for example, by occluding a ribosome binding site, in the case of translation regulation, or forming an intrinsic terminator hairpin, in the case of transcription regulation. Interaction between the sense target and antisense RNAs can then cause structural rearrangement, ultimately controlling the expression of the gene. Years of research have uncovered design principles for these mechanisms, enabling engineers to create a wide array of sRNA regulators, including RNA degradation controllers (Carothers et al., 2011; Carrier and Keasling, 1999), translational repressors (Mutalik et al., 2012; Na et al., 2013) translational activators (Green et al., 2014; Isaacs et al., 2004; Rodrigo et al., 2012), and transcriptional repressors (Takahashi and Lucks, 2013).

Despite the versatility of engineered sRNA regulators, until recently there were no known natural or synthetic examples of sRNAs that could activate transcription (Chappell et al., 2013). To address this gap, we created small transcription activating RNAs (STARs) (Chappell et al., 2015) (Fig. 1). In the STAR mechanism, the sense target region contains an intrinsic terminator hairpin, which terminates transcription in the OFF state, preventing read-through of the downstream gene. The STAR antisense contains a specific anti-terminator sequence that is designed to bind to the 5’ stem of the terminator in *trans* to prevent terminator formation and allow transcriptional read-through in the ON state. This mechanistic design strategy was applied to target a range of intrinsic terminators derived from natural sources, ranging from the pbuE riboswitch to the pT181 plasmid copy control element, ultimately creating five different STARs that displayed a range of transcriptional activation from 3-fold to 94-fold (Chappell et al., 2015). In addition, orthogonality between these STARs and a preexisting library of RNA transcriptional repressors (Takahashi and Lucks, 2013) allowed the construction of two previously unattainable RNA-only logic gates, demonstrating the potential of STARs for engineering sophisticated RNA genetic circuitry.

**Figure 1.**
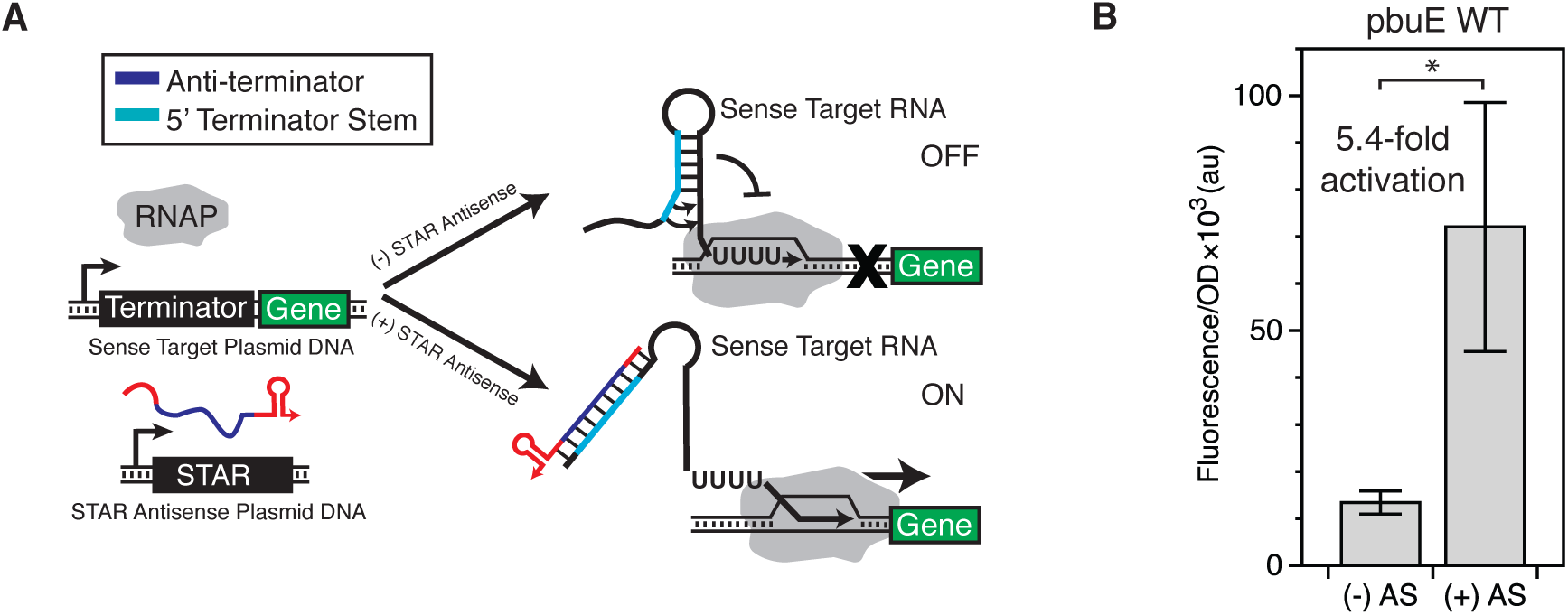
Design and function of a model Small Transcription Activating RNA (STAR). **(A)** Schematic of the mechanism following Chappell et al. (2015). In the absence of the STAR antisense RNA, the nascent sense target RNA upstream of the reporter gene forms a terminator hairpin that stops transcription before the gene is transcribed (OFF). The STAR antisense is designed to contain an anti-terminator sequence complementary to the 5’ side of the terminator hairpin. When the STAR antisense is present, it binds to the terminator sequence, preventing formation of the terminator hairpin and allowing transcription of the downstream gene (ON). **(B)** *In vivo* characterization of the pbuE STAR regulator using superfolder GFP (SFGFP) fluorescence to measure gene expression from a sense target plasmid with and without a STAR antisense (AS) plasmid. Normalized fluorescence was divided by OD 600 to give FL/OD, and fold activation was calculated as FL/OD ON divided by FL/OD OFF. Error bars represent sample standard deviation over 3 independent replicates with 3 colonies each (n=9). The * symbol indicates a statistically significant (p<0.05) increase in FL/OD in the case with STAR antisense as determined by a two-sided T-test.

STARs thus represent a powerful expansion of the RNA engineering toolbox for precisely regulating gene expression and creating synthetic genetic networks. However, much work remains to be done to broadly optimize the fold activation of these new regulators. In particular, only two of the originally designed STARs showed levels of activation greater than 10-fold, thereby limiting the number of STARs useful in applications that require large differences between ON and OFF expression states. Here we remedy this by applying several sRNA engineering methods (Carrier and Keasling, 1997; Sakai et al., 2013) and gene expression optimization strategies to optimize the overall fold activation of weak STAR regulators. In particular, we used a combination of promoter strength tuning and STAR antisense RNA stabilization strategies to improve fold activation from 5.4-fold (±2.2) to 13.4-fold (±3.8) for the pbuE STAR regulator. To confirm that our approach could be generalized, we then applied these strategies to the unrelated prgX STAR (derived from a conjugation control system terminator) to improve its fold activation from 2.1-fold (±0.4) to 14.6-fold (±3.7). This process also yielded multiple STAR variants for both systems with intermediate fold activation levels, showing that this strategy can be used to fine-tune STAR performance. Finally, although the optimization strategies required the addition of a significant amount of extra RNA sequence and structure, orthogonality was preserved between the optimized STARs and a set of antisense RNA transcriptional repressors. These optimization strategies open the door for creating a generation of additional STARs for use in a broad array of biotechnologies.

## Materials and Methods

### Plasmid construction and cloning

STAR-mediated gene expression was tested with a two-plasmid system. Plasmids were constructed so that the sequence encoding each sense target RNA was placed downstream of a constitutive promoter and upstream of the coding sequence for the superfolder green fluorescent protein (SFGFP) reporter (Pédelacq et al., 2006), complete with its own ribosome binding site (Fig. S1). Separate plasmids were constructed for STAR antisense expression, with the sequence encoding the STAR preceded by a constitutive promoter and followed by the t500 transcriptional terminator (Yarnell and Roberts, 1999) (Fig. S1). For experiments in which the STAR antisense is absent, a control plasmid was constructed containing the constitutive promoter followed directly by a transcriptional terminator (rrnB terminator ‘TrrnB’) (Fig. S1).

All plasmids and sequences used in this study are enumerated in Table SI. All sense target plasmids included the p15A origin and a gene for chloramphenicol resistance, while all STAR antisense plasmids contained the ColE1 origin and encoded a gene for carbenicillin resistance. All plasmids were either previously reported or constructed from previously reported plasmids (Chappell et al., 2015; Takahashi and Lucks, 2013) using inverse PCR (iPCR) to make substitutions and/or insertions. The inserted sRNA scaffold and stability hairpin sequences were derived from previous work by Sakai et al. (2013) and Carrier et al. (1999). Sequence-verified stocks of the plasmids were used for all experiments.

### Strains, media, and in vivo bulk fluorescence experiments

All bulk fluorescence experiments were performed in *Escherichia coli* (*E. coli*) strain TG1 with three independent replicates, except for the *hfq* knockout experiments (Fig. 3D), which were performed in *E. coli* strain BW25113 and the BW25113 *–hfq* variant from the Keio collection (Baba et al., 2006). For each independent replicate, pairs of sense target plasmid and STAR antisense (or no antisense control) plasmid were transformed into chemically competent *E. coli* TG1 cells, plated on Difco LB + Agar plates with 100 mg/mL carbenicillin and 34 mg/ml chloramphenicol, and incubated overnight at 37 **°**C for approximately 17 hours. For every experiment the same procedure was repeated for a growth control strain containing plasmids JBL001 and JBL002. Next, the plates were removed from the incubator and kept at room temperature for approximately 7 hours. Three separate colonies were picked for each condition/control, and each was used to inoculate 300 μL of LB media with 100mg/mL carbenicillin and 34 mg/mL chloramphenicol in a 2 mL 96-well block (Costar 3960). In the case of the pbuE variants in Figure 4A, 9 colonies were picked in order to guarantee enough passed the growth requirements described below. The block was covered with a breathable seal (Aeraseal BS-25) and incubated at 37 **°**C while shaking at a speed of 1,000 rpm in a Labnet Vortemp 56 bench-top shaker for 18-19 hours overnight. From this overnight culture 4 μL were taken and used to inoculate 196 μL of M9 minimal media (1 × M9 minimal salts, 1 mM thiamine hydrochloride, 0.4 % glycerol, 0.2 % casamino acids, 2 mM MgSO4, 0.1 mM CaCl2) containing 100 mg/mL carbenicillin and 34 mg/ml chloramphenicol. The cultures, alongside three wells of an M9 only control, were grown in a 2 mL 96-well block under the same conditions as the overnight culture, until the majority of the OD values exceeded 0.07, which took 5 to 7 hours. Fifty μL of this culture were diluted 1:1 with phosphate buffered saline (PBS) in a black-welled clear-bottomed 96-well plate (Costar 3231). The diluted cultures’ optical density (OD) at 600 nm and fluorescence (485 nm excitation, 520 nm emission) were then measured with a Biotek SynergyH1 plate reader.

### Data analysis for bulk fluorescence experiments

Each experiment included two sets of controls: three wells of a media blank (M9) and three wells inoculated from separate colonies the growth control of *E. coli* cells lacking SFGFP but harboring plasmids with the same backbones and resistances as all sense target and STAR antisense plasmids (transformed control JBL001 and JBL002). All fluorescence and OD values for each colony were initially corrected by subtracting the corresponding values from the average of the three media blanks. The ratio of fluorescence units over OD (FL/OD) was then calculated for each well and corrected for background fluorescence by subtracting the average FL/OD for the growth control of cells without SFGFP. For each STAR-target pair, three independent colonies were characterized from three independent transformations (9 colonies total). Data were discarded for colonies that showed low growth (OD<0.07), although this requirement was relaxed for the orthogonality grid in Figure 5 in order to account for the different growth rates of the tested variants. Averages and standard deviations (depicted by error bars) of FL/OD were calculated over the repeat experiments. Fold activation was calculated by dividing the average corrected FL/OD for a STAR-target pair by the average corrected FL/OD for the same target sense plasmid paired with the control no STAR antisense plasmid (JBL002). Fold activation error was calculated using standard error propagation formulas based on the standard deviations of the average corrected FL/OD values. In calculating fold repression (Figure 5), the negative reciprocal was taken to give the fold repression, i.e., 0.20 became -5-fold repression (Chappell et al., 2015).

## Results and Discussion

### The pbuE STAR as a case study for optimization of fold activation

As a starting point for exploring strategies to increase fold activation, we chose to focus on a STAR-target system (Chappell et al., 2015) derived from the intrinsic terminator of the pbuE riboswitch (Ceres et al., 2013b), which showed a low fold activation of transcription (Figure 1B). As illustrated in Figure 1A, the pbuE STAR is designed to be fully complementary to the 5’ half of the pbuE intrinsic terminator present in the target RNA. Interaction of the STAR with its target RNA thus prevents the formation of the terminator hairpin, enabling transcription elongation into the downstream reporter gene, superfolder green fluorescent protein (SFGFP). To determine fold activation, gene expression was characterized by measuring fluorescence normalized by optical density (FL/OD) for cultures of *E. coli* cells co-transformed with two plasmids: one plasmid encoding the pbuE sense target fused to the downstream SFGFP coding sequence, and the other plasmid encoding either the STAR antisense (ON state) or an empty backbone control (OFF state) (absence of STAR antisense case). Activation was then calculated as a ratio of the ON/OFF FL/OD values (see Materials and Methods). For the original pbuE wild-type STAR (WT), we observed 5.4-fold (±2.2) activation in the presence of the STAR antisense compared to when only the target RNA was expressed (Fig. 1B). This result indicated that there was ample room for improvement in fold activation compared to the 94-fold (±26) activation shown by the best STAR activator reported previously (Chappell et al., 2015). Furthermore, previous work on applying RNA sequence optimization strategies did not improve the low fold activation of the pbuE STAR system (Chappell et al., 2015), motivating us to pursue a suite of alternative strategies discussed below.

### Improving fold activation of STARs by manipulating STAR/target expression ratios

To begin, we chose to investigate the possibility of improving fold activation by increasing the relative concentration ratio of STAR antisense to its complementary sense target. A previously developed model of the STAR mechanism hypothesized that transcription activation is directly related to the rate of binding between STAR and target (Chappell et al., 2015). Therefore, increasing the expression level of the STAR antisense relative to its target should naturally increase the number of binding events, and thus increase the likelihood of any given target RNA being transcriptionally activated. To test this, we manipulated the STAR/target expression ratio *in vivo* in *E. coli* by altering the relative strengths of the constitutive promoters that drive the expression of the STAR antisense and target sense in our two-plasmid system (see Materials and Methods). Given that both the STAR and target RNAs were originally under the control of the same strong constitutive σ^70^ promoter, decreasing the strength of the target RNA promoter provided a straightforward way to titrate down the steady-state levels of sense target RNA by reducing the transcription rate. Following this strategy, we cloned a series of successively weaker constitutive promoters upstream of the pbuE target sense RNA and examined their effects on fold activation *in vivo* (Fig. 2). We chose promoters from the Anderson promoter library from the Registry of Standard Biological Parts (partsregistry.org), whose strengths have been well-characterized in previous work (Kelly et al., 2009) (Table SII).

**Figure 2.**
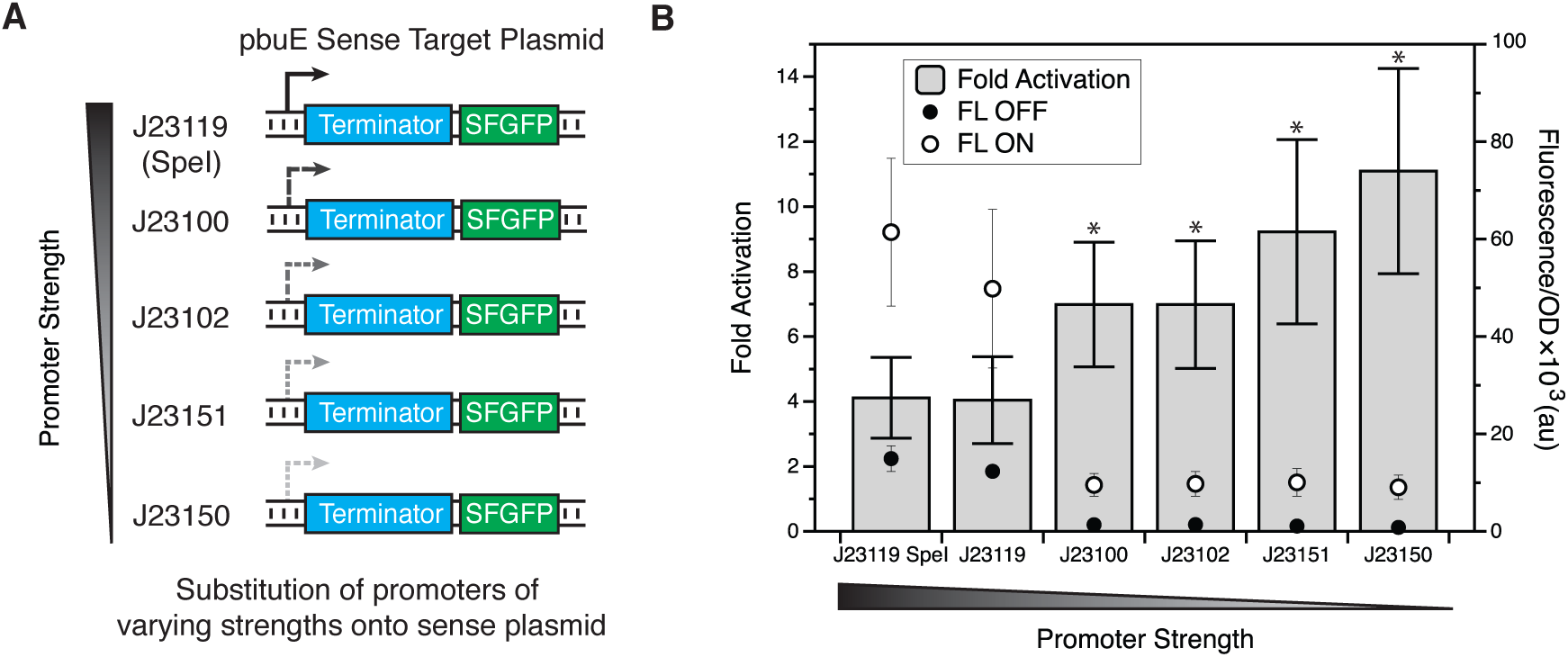
Optimization of the STAR/target expression ratio yields higher fold activation.**(A)** Schematic of the design strategy for improved activation. A number of weaker promoters were substituted for the strong J23119 (SpeI) promoter on the target sense plasmid in order to decrease the expression level of the sense RNA. In these experiments, the STAR was expressed from a high-copy plasmid using the strong J23119 (SpeI) promoter. This promoter series was designed to increase the relative expression ratio of STAR to target. **(B)** *In vivo* fluorescence characterization demonstrates an increase in fold activation when a weaker target promoter is used, with a fold activation of 11.1-fold (±3.6) observed for the weakest J23150 promoter. Normalized fluorescence divided by OD 600 (FL/OD) is plotted against the right-hand axis as a series of circles, with the closed circles representing the FL OFF level (no STAR control plasmid) and the open circles representing the FL ON level (STAR present). Fold activation (FL/OD ON divided by FL/OD OFF) is plotted against the left-hand axis as a series of gray bars. Error bars represent sample standard deviation over 3 independent replicates with 3 colonies each (n=9). The * symbol indicates statistically significant (p<0.05) improvement in ON level (normalized to OFF level) over the wild-type sense promoter configuration (J23119 SpeI) as determined by a two-sided T-test.

**Figure 3.**
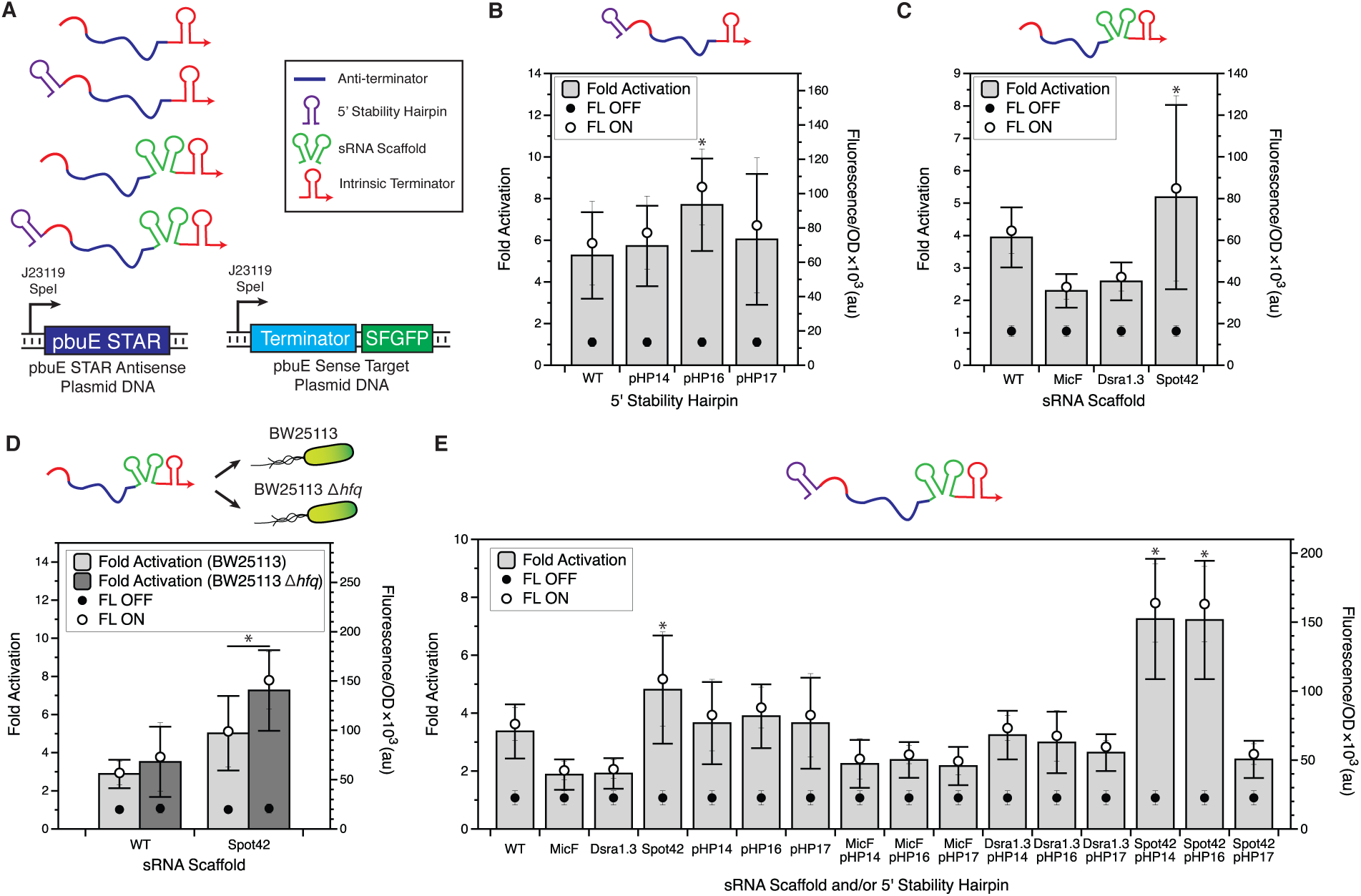
**(A)** Schematic of the changes made to the pbuE STAR to improve stability. Stability hairpins (purple) were added to the 5’ end of the pbuE STAR to block RNase E-mediated degradation (Carrier and Keasling, 1997). sRNA scaffolds (green) were added to the 3’ end to improve sRNA stability through the recruitment of Hfq, a protein important for mediating and stabilizing sRNA interactions (Vogel and Luisi, 2011). **(B)** *In vivo* functional characterization of stability hairpin STAR variants indicates that the addition of 5’ stability hairpins, particularly pHP16, confers a modest increase in activation function to the pbuE STAR regulator. **(C)** *In* vivo functional characterization of sRNA scaffold STAR variants indicates that only the addition of the Spot42 sRNA scaffold results in a small increase in fold activation. Addition of the MicF or Dsra1.3 sRNA scaffolds to the pbuE STAR was shown to decrease fold activation. **(D)** Testing of the Spot42 sRNA scaffold variant of the pbuE STAR in BW25113 and BW25113 *–hfq* demonstrates slightly *higher* activation levels in the absence of Hfq. **(E)** Combining both sRNA scaffolds and 5’ stability hairpins yields the highest fold activation for the combinations of Spot42/pHP14 and Spot42/pHP16. Data is plotted as in Figure 2. Wild-type (WT) indicates the initial pbuE STAR regulator using the strong J21339 (SpeI) promoter to drive both STAR and target RNA expression. In parts **(B)**, **(C)**, and **(E)** the * symbol indicates a statistically significant (p<0.05) increase in ON level FL/OD over WT as determined by a two-sided T-test. In part **(D)** * indicates a statistically significant (p<0.05) difference in ON level (normalized to OFF level) between the two strains tested as determined by a two-sided T-test.

The *in vivo* testing of these target promoter variants indicated that weakening the sense plasmid promoter strength did indeed result in greater fold activation, and we observed a clear correlation between decreased sense promoter strength and increased fold activation (Fig. 2B). Although the overall ON level of fluorescence decreased as expected with a weaker promoter on the target sense plasmid, we observed an even greater decrease in the OFF level, leading to an overall increase in fold activation. When compared to the original promoter configuration, the weakest promoter tested, J23150, more than doubled the fold activation of the pbuE STAR regulator from 4.1-fold (±1.2) to 11.1-fold (±3.2). These results demonstrate that we can increase the fold activation of STAR regulators through manipulating STAR/target expression ratios.

### Improving fold activation of STARs by stabilizing the STAR antisense RNA

Since manipulating RNA stability is a key point of control for RNA mechanisms (Chappell et al., 2013; Smolke and Keasling, 2002), we next sought to examine how altering the stability of the STAR antisense could be used as another strategy for improving fold activation. The steady-state level of STAR antisense RNA molecules available for activation at any time is governed by the balance between two key rates: the rate of synthesis (transcription) and the rate of degradation. Since the STAR antisense was already expressed from a high-copy plasmid (colE1 origin of replication) under the control of a strong promoter, reducing the degradation rate presented a more accessible way to increase the level of STAR antisense RNA and thus fold activation. In order to decrease the degradation rate of the STAR antisense RNA, we sought to use two RNA engineering strategies: (i) stabilizing the STAR by adding strong RNA hairpins to the 5’ ends, a strategy that has been shown in previous work to stabilize mRNAs (Carrier and Keasling, 1999), and (ii) adding a naturally occurring sRNA scaffold to the 3’ end, which has been recently demonstrated to improve the function of small translational and transcriptional RNA regulators (Sakai et al., 2013) (Fig. 3A).

Our first strategy for stabilizing antisense RNAs was based on the fact that secondary structures located at the 5’ end of bacterial mRNAs can confer stability (Carrier and Keasling, 1997; Emory et al., 1992) by blocking RNAse E-mediated degradation. In particular, this strategy has been shown to improve mRNA stability and lengthen mRNA half-lives (Carrier and Keasling, 1999). Moreover, a strong correlation has been observed between the computationally predicted secondary structure free energies of these 5’ hairpins and the steady-state levels of mRNA, allowing for the design of RNA hairpins that confer varying levels of stability (Carrier and Keasling, 1999). To investigate whether or not these stability hairpins added to STARs would improve fold activation, we added three previously published synthetic RNA stability hairpins (pHP14, pHP16, and pHP17) (Carrier and Keasling, 1999) to the 5’ end of the pbuE STAR. Before cloning these fusions, we first used computational RNA structure modeling with RNAStructure (Reuter and Mathews, 2010) to confirm that the hairpins were not predicted to interfere with the folding of the pbuE STAR antisense (Fig. S2). We then cloned the hairpins onto the 5’ end of the pbuE STAR sequence, directly after the promoter, and characterized the resulting fold activation of these modified STARs (Fig. 3B). Compared to the WT pbuE STAR, one of the variants with the added 5’ stability hairpin demonstrated modest improvements in *in vivo* fold activation (Fig. 3B). In particular, we observed that fold activation was increased from 5.2-fold (±2.1) to 7.7-fold (±2.2) with pHP16. Interestingly, the level of increased activation did not directly correlate with the previously reported half-lives of these stability hairpins (Carrier and Keasling, 1999). While pHP14, pHP16, and pHP17 had been shown to grant successively longer half-lives (in numerical order), in the context of the pbuE STAR, only pHP16 appeared to confer a statistically significant increase in fold activation. This result could indicate that stability gained by the addition of a particular hairpin may be governed by the local sequence context or that other structural effects are causing these differences in fold activation.

As an alternate method of increasing fold-activation through stabilizing the STAR antisense, we tested adding sRNA-derived scaffolds to the 3’ end of the STAR, upstream of the transcriptional terminator. This strategy was based upon recent research by Sakai et al. (2013) showing that the addition of sRNA scaffolds to translational sRNA activators can lead to higher fold activation. Not only do the scaffolds stabilize the RNA through the addition of secondary structure, these particular sRNA-derived scaffolds are also designed to include binding sites for the RNA-binding chaperone protein Hfq (Sakai et al., 2013). Among its many roles, Hfq is known to aid sRNA function by mediating sRNA interactions with target mRNAs for many trans-encoded sRNAs that regulate translation (Møller et al., 2002; Zhang et al., 2003). Hfq can also modulate sRNA stability by affecting ribonuclease accessibility or susceptibility to 3’ polyadenylation and subsequent degradation (Vogel and Luisi, 2011). To test the ability of sRNA scaffolds to increase fold activation for STAR regulators, we chose the MicF, Dsra1.3 and Spot42 scaffolds that were previously shown to lead to the greatest improvements in activation and repression levels in the RNA regulators tested (Sakai et al., 2013). We then fused each scaffold to the 3’ end of the pbuE STAR, directly before the transcriptional terminator (Fig. 3A). Functional testing *in vivo* showed variable results depending on the scaffold used. In particular, while the addition the Spot42 scaffold to the pbuE STAR antisense slightly increased activation, both MicF and Dsra1.3 markedly decreased activation levels (Fig. 3C). Structural prediction with RNAStructure (Reuter and Mathews, 2010) indicated that a number of the most probable low-energy structures for the antisense RNA fused with MicF or Dsra1.3 interfered with the native structure of both the STAR antisense and the sRNA scaffolds (Fig. S3). Thus the observed decrease in activation could be the result of structural interference between the scaffold and the STAR.

We next sought to test the role of Hfq in the observed increase in STAR activation with the Spot42 scaffold. To test this, we repeated the *in vivo* functional characterization of the Spot42 fusions in both the Keio collection Δ*hfq* knockout strain and its parent *E. coli* K-12 BW25113 strain (Baba et al., 2006) (Fig. 3D). We found that the absence of Hfq had little to no effect on the wild-type pbuE STAR activation, as expected given that the wild-type does not contain an Hfq-recruiting scaffold sequence. Surprisingly, we found that the activation level of the pbuE STAR-Spot42 fusion significantly improved in the absence of Hfq, in contrast to previous observations of the reliance of sRNA scaffolds on Hfq for added stability (Sakai et al., 2013). One possible explanation for this effect is that over-expression of a small RNA with a high-affinity Hfq binding site interferes with basic cellular processes by creating competition for Hfq, an effect that has been observed previously (Moon and Gottesman, 2011).

Next, we examined whether we could increase fold activation further by combining the stability hairpin and sRNA scaffold strategies together. To test this, we created all possible combinations of the pbuE STAR antisense containing both 5’ stability hairpins and 3’ sRNA scaffolds. *In vivo* characterization indicated that the combination of the Spot42 scaffold with either the pHP14 or pHP16 stability hairpin granted improved activation over any of the variants with only hairpin or scaffold (Fig. 3E). In particular, we observed 7.2-fold (±2.1) activation for the variant with both Spot42 and pHP14 and 7.2-fold (±2.0) activation for the variant with Spot42 and pHP16, both better than the 4.8-fold (±1.9) activation seen for Spot42 alone.

Interestingly, these increases in fold activation are larger than the improvements in transcription repression seen when scaffolds were applied to the pT181 RNA-based transcriptional repressor (Sakai et al., 2013). However, Sakai et al. were able to successfully use these strategies to improve the fold activation of an RNA translational activator. These results could suggest a more general principle for antisense RNA stabilization strategies being more effective for gene expression activation, or could indicate these strategies are more effective when applied to relatively unstructured antisense RNAs.

Overall, our results showed that the incremental improvements in fold activation generated by the addition of 5’ stability hairpins and 3’ scaffolds alone can be combined in a modular and synergistic fashion to generate STARs with even higher fold activation.

### A general method for increasing STAR activation by combining expression level tuning and RNA stability strategies

We subsequently investigated whether combining expression level tuning and RNA stabilization strategies could be used to further optimize fold activation for the pbuE STAR. To do this, we combined the Spot42/pHP14 pbuE antisense variant together with the weakened promoter strength series on the target sense RNA. Combining the stabilized STARs with the weaker promoter target sense plasmids yielded another increase in activation level, reaching as high 13.4-fold (±3.8) activation in the case of the Spot42 pHP14 antisense combined with the target sense plasmid containing the J23151 promoter (Fig. 4A). While the general trend was toward higher fold activation with weaker promoters, there was high error and the trend was less uniform than seen previously. Nevertheless, the higher fold activation seen from combining the optimized pbuE STAR with weaker target sense promoters indicates the modularity of these two independent strategies for increasing fold activation.

**Figure 4.**
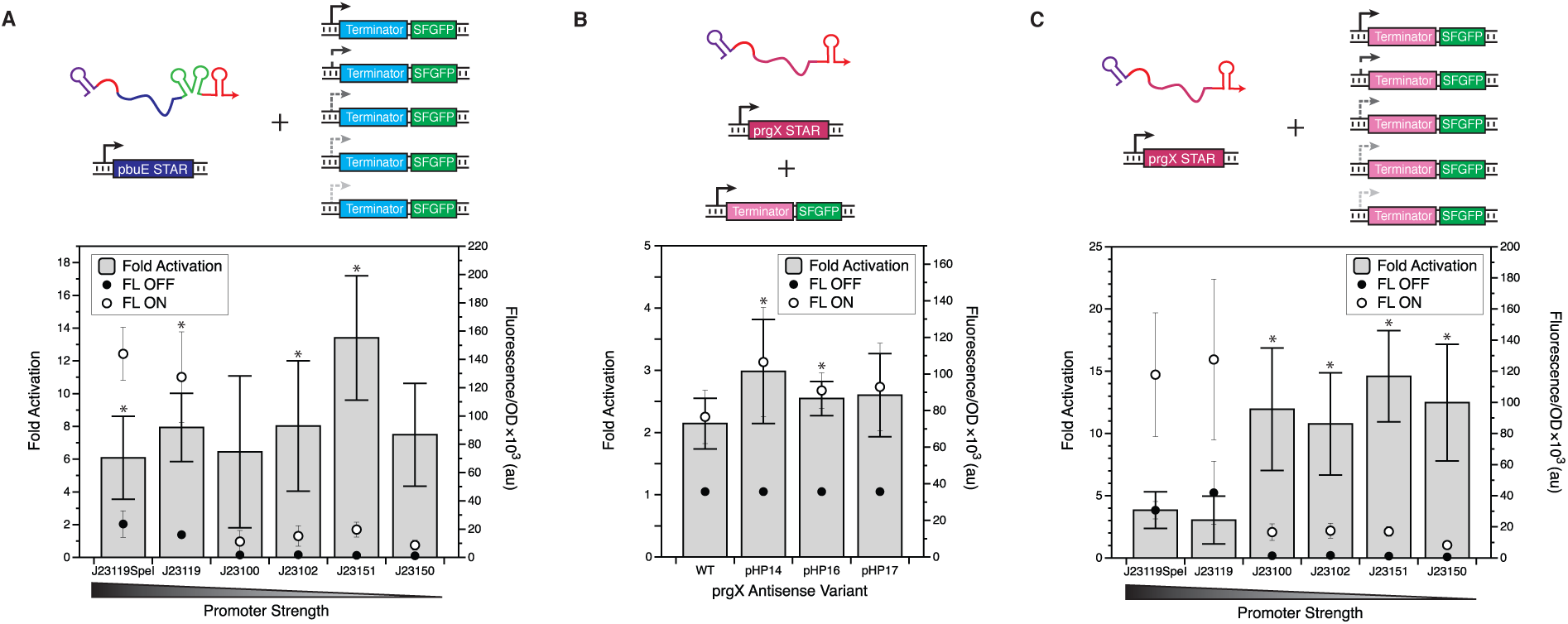
Combining expression level tuning and RNA stability strategies improves STAR fold activation in multiple systems. **(A)** Combining the stabilized pbuE STAR antisense (Spot42/pHP16) with weaker target sense promoters yields up to 13.4-fold (±2.0) activation for the J23151 promoter pairing. **(B)** 5’ stability hairpins slightly improve fold activation for the prgX STAR regulator. **(C)** Combining the prgX STAR stabilized by pHP14 with weaker strength target promoters yields another boost in fold activation, showing the broader applicability of these STAR optimization strategies. Data is plotted as in Figure 2. Wild-type (WT) indicates the initial pbuE STAR regulator in part **(A)** and the initial prgX STAR regulator in part **(B)** and **(C)**, both of which use the strong J23119 (SpeI) promoter to drive both STAR and target RNAs. In parts **(A)** and **(C)** * indicates a statistically significant (p<0.05) increase in ON level (normalized to OFF level) over WT, as determined by a two-sided T-test. In part **(B)** the * symbol indicates a statistically significant (p<0.05) increase in ON level FL/OD over WT as determined by a two-sided T-test.

Having successfully optimized the pbuE transcriptional activator from its initial 5.3-fold (±2.2) activation level to 13.4-fold (±2.0) activation, we next sought to test the generality of this combined method for improving STAR fold activation. We started with a previously constructed STAR generated from the prgX conjugation control system (Weaver, 2007) that only displayed a 2.1-fold (±0.4) activation (Chappell et al., 2015) (Fig. 4B). In particular, we applied the same 5’ stability hairpins as used above and the Spot42 sRNA scaffold to the prgX STAR. *In* vivo testing revealed that while some of the hairpin additions modestly improved fold activation (Fig. 4B), the addition of the Spot42 scaffold did not (Fig. S4). When combined with different strength sense target promoters, the best STAR variant (pHP14) had an even greater increase in fold activation (Fig. 4C), providing further proof of the modularity of the strategies for modifying sense promoter and STAR antisense stability. Overall, the best prgX variant displayed 14.6-fold (±3.7) activation, a vast improvement from the original 2.1-fold (±0.5) activation and a validation that the optimization strategies work on an additional, unrelated STAR system.

### Testing the orthogonality of optimized STARs to themselves and to RNA transcriptional repressors

We next sought to test whether our two newly optimized STARs were orthogonal to the previously reported ribA STAR and three previously developed transcriptional repressors (Takahashi and Lucks, 2013). If two STARs are orthogonal, then the STAR antisense of one should not regulate the target sense RNA of the other (and vice versa), allowing them to be used together in a complex regulatory system without cross-talk. Such orthogonality is non-trivial, especially given that the addition of additional RNA sequence to the optimized STARs increases the potential for off-target interactions, as does increasing the concentration ratios of STARs to their targets. Moreover, orthogonality of these component parts is key for their use in the higher-order logic gates and circuits that STARs and transcriptional repressors have been used to construct (Chappell et al., 2015; Takahashi and Lucks, 2013). For example, the ability to build circuits using orthogonal elements allows synthetic biologists to program systems with complex functionalities like ligand-sensitive NOR gates (Qi et al., 2012) and RNA cascades that control the timing of gene expression (Takahashi et al., 2014), a vital capability for biotechnology applications.

To perform the orthogonality test, we challenged each optimized STAR antisense variant against the best target senses from the optimized pbuE and prgX systems (using the J23151 promoter), along with the target senses from the previously reported ribA STAR (Chappell et al., 2013). We also checked for orthogonality to target regions from three RNA transcriptional repressors: pT181.H1, Fusion 4, and Fusion 6 from Takahashi and Lucks (2013). As a further test, we challenged each RNA transcriptional repressor antisense variant against the same set of target senses. This resulted in a 6x6 matrix of conditions demonstrating the observed orthogonality of these regulators to one another (Fig. 5, Fig. S5). Both of the newly optimized STARs showed reasonable orthogonality to the other regulators. Though the pbuE STAR antisense did exhibit slight cross-talk with the prgX target sense, the effect was still well below the activation seen with the cognate STAR antisense.

**Figure 5.**
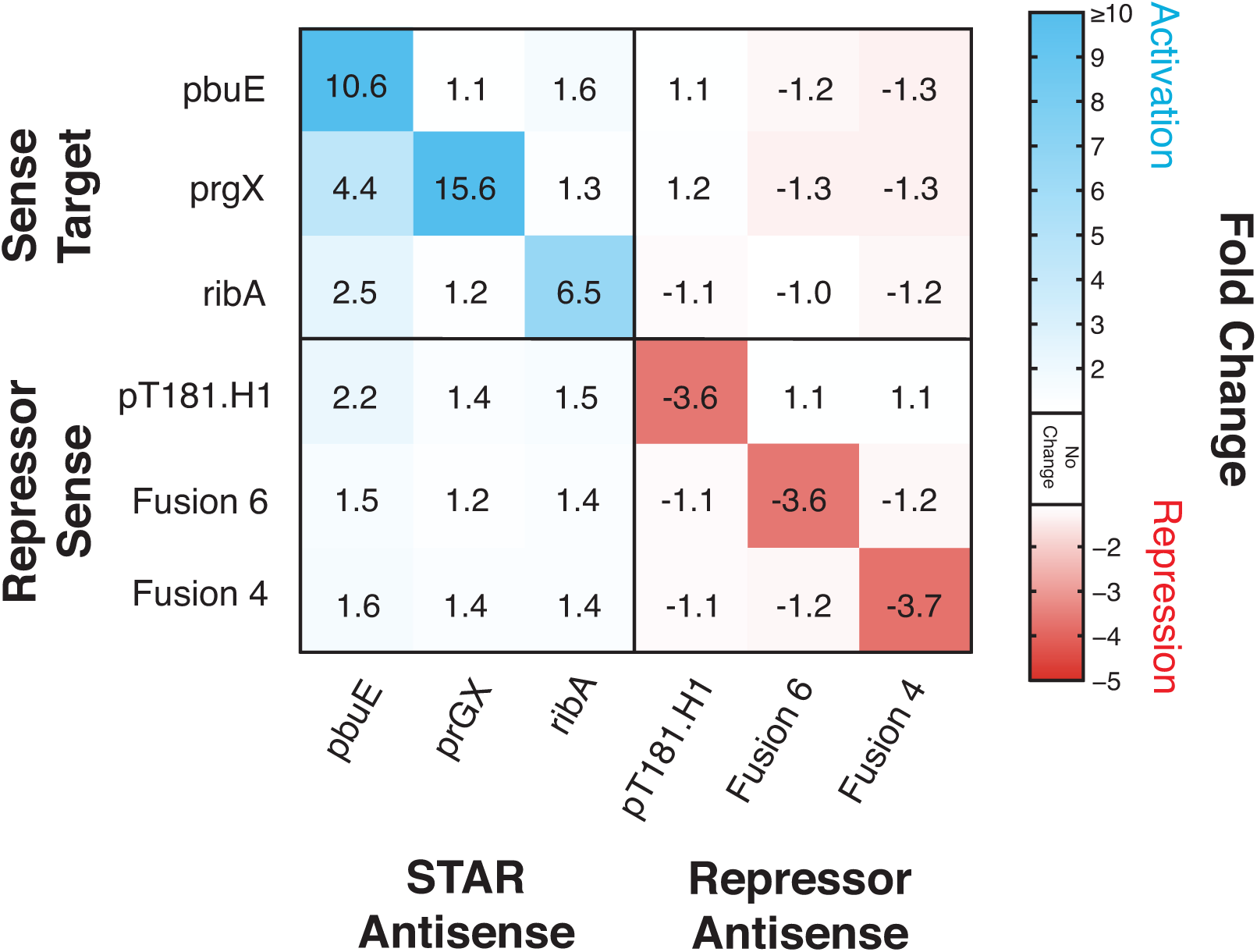
Testing the orthogonality of improved STAR regulators with themselves and with RNA transcriptional repressors. Characterization of a 6 × 6 orthogonality matrix consisting of the newly optimized pbuE and prgX STAR regulators, the previously reported ribA STAR regulator (Chappell et al., 2015), and the pT181.H1, Fusion 4, and Fusion 6 transcriptional repressors reported by Takahashi and Lucks (2013). Each matrix square represents the fold change of gene expression for the indicated combination of STAR or repressor plasmid and target plasmid, compared to the condition with target plasmid and an empty control plasmid. Fluorescence characterization (measured in units of FL/OD, fluorescence divided by OD at 600 nm) was used to calculate average fold change, which is represented by a color scale in which ≥ 10-fold is the darkest blue (activation), 1-fold is white (no activation or repression) and -5-fold is red (repression). FL/OD plots for each individual combination are shown in Fig. S5. Data represents mean values of *n* = 9 biological replicates.

These results confirm that these STAR optimization strategies largely do not affect the orthogonality of STARs between themselves and RNA transcriptional repressors. Not only will the addition of new orthogonal STARs allow for more complex RNA circuitry, the optimization strategies used to improve STAR activation will allow for the future development of more highly functional RNA transcriptional activators.

## Conclusions

In this work, we tested several RNA engineering strategies for optimizing the fold activation of small transcription activating RNAs. In particular, we focused on strategies designed to stabilize the STAR antisense and alter the concentration ratio between the STAR and its target. Using the pbuE STAR as a test case, we showed that the addition of 5’ stability hairpins and scaffolds to the STAR antisense and the ability to adjust the ratio of STAR antisense to target sense via promoter strength tuning give convenient, modular, and synergistic ways to alter the transcription activation levels of two distinct STAR systems. Specifically, we found that these strategies as applied to the pbuE STAR system increased the fold activation from 5.3-fold (±2.2) to 13.4-fold (±2.0). Moreover, we showed that they were general, and when applied to the unrelated prgX STAR regulator, yielded an increase in transcription activation from 2.1-fold (±0.4) to 14.6-fold (±3.7). Furthermore, we showed that these changes largely preserved the orthogonality of these optimized STARs to themselves and to a panel of RNA transcriptional repressors that have been used to construct higher-order RNA transcriptional circuits.

These results are significant for several reasons. First, these optimizations have expanded the repertoire of STARs - the starting point fold activations for the pbuE and prgX STAR regulators prohibited their use for higher order circuit construction, a problem remedied by our optimizations. Second, the optimization strategies used on the pbuE and prgX systems should be applicable to many other STARs, paving the way for even larger libraries of RNA regulators. Finally, the demonstrated orthogonality between the newly optimized activators and previously reported regulators, in addition to representing a non-trivial achievement, makes them highly useful for future circuit-building, allowing for complex genetic logic gates and circuits to be built without interference between different components.

It should be noted that the majority of the improvement in STAR fold activation was achieved through decreasing the OFF level. If a high ON level were required for a particular application, other strategies could be used to increase ON levels while maintaining low OFF levels. For instance, if high protein expression were desired, modular strategies aimed at tuning translation through altering ribosome binding site strength and accessibility could be used as an alternate way of manipulating target gene expression levels.

In addition to optimizing STAR fold activation, the tested strategies have led to a series of STARs with varying ON, OFF and fold activation levels. These strategies have thus created a panel of variants that can be used for fine-tuning of transcription activation. The importance of fine-tuning individual regulators to enable the correct performance of a larger circuit has been demonstrated in numerous examples of synthetic circuits (Ellis et al., 2009; Elowitz and Leibler, 2000; Wang et al., 2009), making the suite of functional STAR variants a useful library to draw from for future circuit design. Strategies like these are becoming more important as synthetic biology looks to implement increasingly sophisticated genetic circuitry, requiring the ability to carefully calibrate the function of biological circuit components.

In summary, we have successfully expanded our capabilities for genetic regulation and made highly useful additions to the synthetic biology toolkit through systematic optimizations of a set of small transcription activating RNAs. These newly-improved STAR regulators will allow for the construction of complex cellular circuitry, and the optimization strategies will be highly useful for creating a generation of additional STARs for use in a broad range of biotechnologies.

## Acknowledgements

We thank Professor Matthew DeLisa (Cornell Chemical and Biomolecular Engineering) for providing the BW25113 and BW25113 Δhfq strains. This material is based upon work supported by the National Science Foundation Graduate Research Fellowship Program [DGE-1144153 to S.M.], Defense Advanced Research Projects Agency Young Faculty Award (DARPA YFA) [N66001-12-1-4254 to J.B.L.], and an Office of Naval Research Young Investigators Program Award (ONR YIP) [N00014-13-1-0531 to J. B. L.]. J.B.L. is an Alfred P. Sloan Research Fellow.

The authors declare competing financial interest. The authors have submitted a provisional patent application (No. 61/981,241) for the technologically important developments included in this article.

This is the pre-peer reviewed version of the following article: S. Meyer, J. Chappell, S. Sankar, R. Chew, J. B. Lucks. “Improving fold activation of small transcription activating RNAs (STARs) with rational RNA engineering strategies.” Biotechnology and Bioengineering, 2015, which has been published in final form at http://onlinelibrary.wiley.com/journal/10.1002/(ISSN)1097-0290.

